# Peak performance singing requires daily vocal exercise in songbirds

**DOI:** 10.1101/2023.02.23.529633

**Authors:** Iris Adam, Katharina Riebel, Per Stål, Neil B. Wood, Michael J. Previs, Coen P.H. Elemans

## Abstract

Vocal signals mediate much of human and non-human communication. Key performance traits - such as repertoire size, speed and accuracy of delivery - affect communication efficacy in fitness-decisive contexts such as mate choice and resource competition^1^. Specialized fast vocal muscles^2,3^ are central to accurate sound production^4^, but it is unknown whether vocal, like limb muscles^5,6^, need exercise to gain and maintain peak performance^7,8^. Here, we show that for song development in juvenile songbirds, the closest analogue to human speech acquisition^9^, regular vocal muscle exercise is crucial to achieve adult peak muscle performance. Furthermore, adult vocal muscle performance reduces within two days of abolishing exercise, leading to downregulation of critical proteins transforming fast to slower muscle fibre types. Daily vocal exercise is thus required to both gain and maintain peak vocal muscle performance, and if absent changes vocal output. We show that conspecifics can detect these acoustic changes and females prefer the song of exercised males. Song thus contains information on recent exercise status of the sender. Daily investment in vocal exercise to maintain peak performance is an unrecognized cost of singing and could explain why many birds sing daily even under adverse conditions^10^. Because neural regulation of syringeal and laryngeal muscle plasticity is equivalent, vocal output may reflect recent exercise status in all vocalizing vertebrates.

---

The production of complex, learned vocalisations, such as human speech and birdsong, comprise some of the most complicated movements of the vertebrate body. Precise motor control and execution of these vocalisations plays a decisive role in mate choice and resource competition. Birds^11,12^ and mammals^4,13^ control their vocal organs with highly specialized muscles optimizing for speed. These muscles minimize the duration from neural excitation to muscle contraction compared to limb muscles by multiple molecular and cellular adaptations: *i*) faster calcium signalling by doubling the amount of sarcoplasmic reticulum, *ii*) fast detachment rates by expressing molecular motor isoforms that allow superfast actomyosin crossbridge cycling and *iii*) aerobic, high-energy production by having up to 30% versus 2% of cell volume dedicated to mitochondria^4^ with densely packed cristae^14^. These extreme muscle adaptations suggest critical function in vocal communication systems.

Skeletal muscles demonstrate life-long plasticity in response to changes in functional load or hormonal state, which has been studied extensively in limb muscles for sport science and biomedicine^5,6^. In humans it is unknown if vocal muscles exhibit exercise-induced plasticity in critical features such as speed and force, and if such changes directly affect vocal output^8^. Such insights could aid to provide a mechanistic basis for voice training programs for prevention and performance recovery for voice professionals, the aging voice, and treatment for diseases that affect neuromuscular changes, such as dysarthria and muscle tension dysphonia^8^. However, we lack access to human laryngeal muscular function in vivo, which has heeded the explicit call for animal models that are functionally equivalent to the human voice^8^. Song learning in songbirds is widely accepted as the closest animal analogue for human speech acquisition^9^, and muscles controlling the avian vocal organ, the syrinx, change concurrent with song learning during postnatal development^4,15,16^. Both mass and speed of vocal muscle speed gradually increase over the course of song learning process and with age^4,15,16^. These gradual muscle changes could arise from brain-body interaction via either exercise-induced plasticity^7^ or muscle maturation, but this remains as yet untested.

Many bird species sing daily to tie and maintain social bonds as well as defend territories^17^. However, many birds sing daily outside of these contexts: in captivity, even isolated zebra finches sing hundreds of nondirected songs per day^18,19^, and in the wild birds keep singing daily even under extremely adverse conditions^10^. If birds need to sing daily for exercise to keep their muscles in shape this could explain that they also sing even in situations where their singing would not aid territorial defence, mate attraction or other functions.

Here we test the hypotheses that early acquisition as well as maintenance of adult peak vocal muscle performance requires vocal muscle exercise in a well-studied vocal learner - the zebra finch. We furthermore test if exercise-induced muscle property changes affect vocal production and if conspecifics can detect and evaluate these changes in a mate-choice context.

## Performance acquisition needs exercise

First, to test the hypothesis that exercise is needed to acquire peak adult vocal muscle performance, we exploited the bipartite structure of the songbird syrinx where two bilateral pairs of vocal folds (left and right hemisyrinx) independently contribute to song^20^. We allowed normal muscle use during vocal ontogeny of the left hemisyrinx (i.e., intact side), and experimentally prevented use of the right hemisyrinx by syringeal muscle denervation in juveniles (DPH) (**Fig 1**; See Methods). Next, we measured two key features of physiological muscle performance – duration of muscle twitch contraction, as a proxy for contraction speed, and maximal isometric stress (MIS) in the dorsal tracheosyringeal muscle (DTB). We compared these muscle features on both sides when these animals were adult (100DPH) and song learning was completed. On the intact side, both contraction speed (4.89±0.86 ms, N=7) and MIS (2.61±1.37 mN/mm^2^, N=7) were similar to wildtype adult males (**Fig 1a, b**). However, on the denervated side, contraction speed was halved to 9.13±3.18 ms and MIS quadrupled to 12.23±9.36 mN/mm^2^ (**Fig 1b**), cross-sectional area (CSA) was reduced by 70% (**Fig 1d, e**). The myofiber number remained the same (**Fig 1f**). These large treatment-induced differences in physiological performance were also evident in muscle morphology: The denervated adult hemisyrinx muscle visually and physiologically resembled the juvenile syrinx at 25DPH (**Fig 1c**).

**Figure 1.**
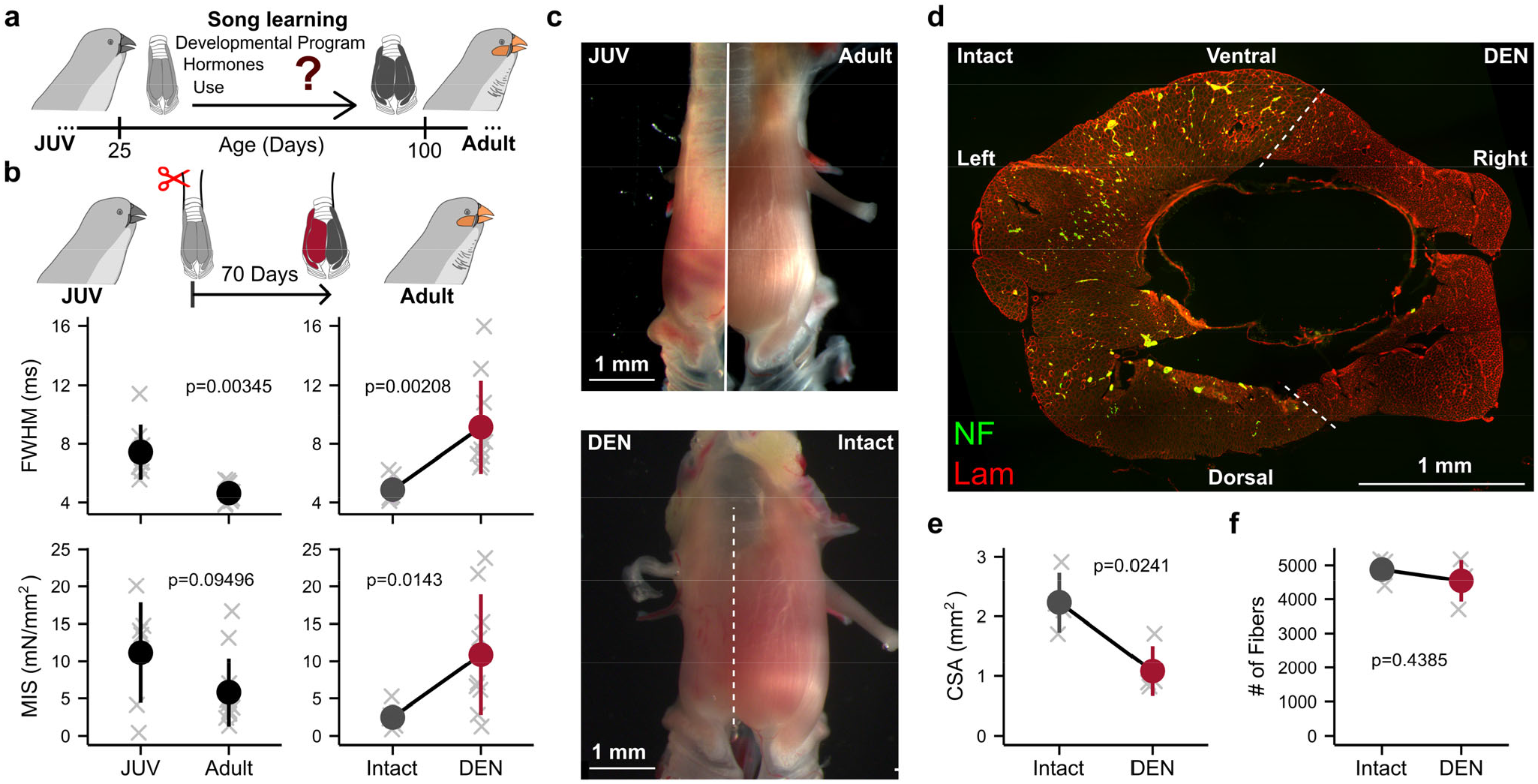
Acquiring adult vocal muscle performance requires exercise during vocal development. a) During song learning vocal muscles in male zebra finches gradually hypertrophy and increase in contraction speed from juvenile (JUV) to adult. b) Denervation (DEN) of the right hemisyrinx at the onset of song learning (red) decreases adult contraction speed, which is measured as the full width at half maximum (FWHM) of isometric force development of a single twitch. Maximal isometric stress (MIS) during tetanic stimulation is also increased compared to the intact side (grey). c) Muscles on the denervated hemisyrinx remain small and resemble juvenile syrinx muscles. d) Immunohistochemical staining of a cross-section of the adult male zebra finch syrinx stained for laminin (Lam, red) to quantify fibre number, cross-sectional area (CSA) and neurofilament (NF) for confirming successful denervation. e) Summed CSA is significantly smaller on the denervated side, while f) the number of muscle fibres remains the same.

Taken together, these data support the hypothesis that achieving adult peak muscle performance requires vocal muscle exercise during the song-learning phase.

## Performance maintenance needs daily exercise

Second, to test whether continued vocal muscle exercise is needed to maintain adult performance, we prevented muscle exercise in adult males by unilateral denervation (**Fig 2a**, see Methods). Muscle speed decreased rapidly (**Fig 2b**) on the denervated side, while it remained unaffected on the intact side. Only two days post-denervation the DTB muscle had slowed down, and after 21 days post-denervation its speed reverted to juvenile levels (**Fig 2b**). The effects of denervation on MIS were even more dramatic: merely two days post denervation MIS dropped fivefold compared to the intact side (from 7.13±4.85 to 1.43±1.88 mN/mm^2^, **Fig 2c**), which remained unaffected compared to wildtype. Morphologically, denervation reduced the CSA to 74±8% (N=3) of the intact side 21 days post-denervation (**Fig 2d**) while the number of muscle fibres remained the same (**Fig 2e**). Thus, denervation severely affects key physiological and morphological muscle features that cause these vocal muscles to lose their peak performance within days. Taken together, these data support the hypothesis that maintaining adult peak muscle performance requires regular, daily vocal muscle exercise.

**Figure 2.**
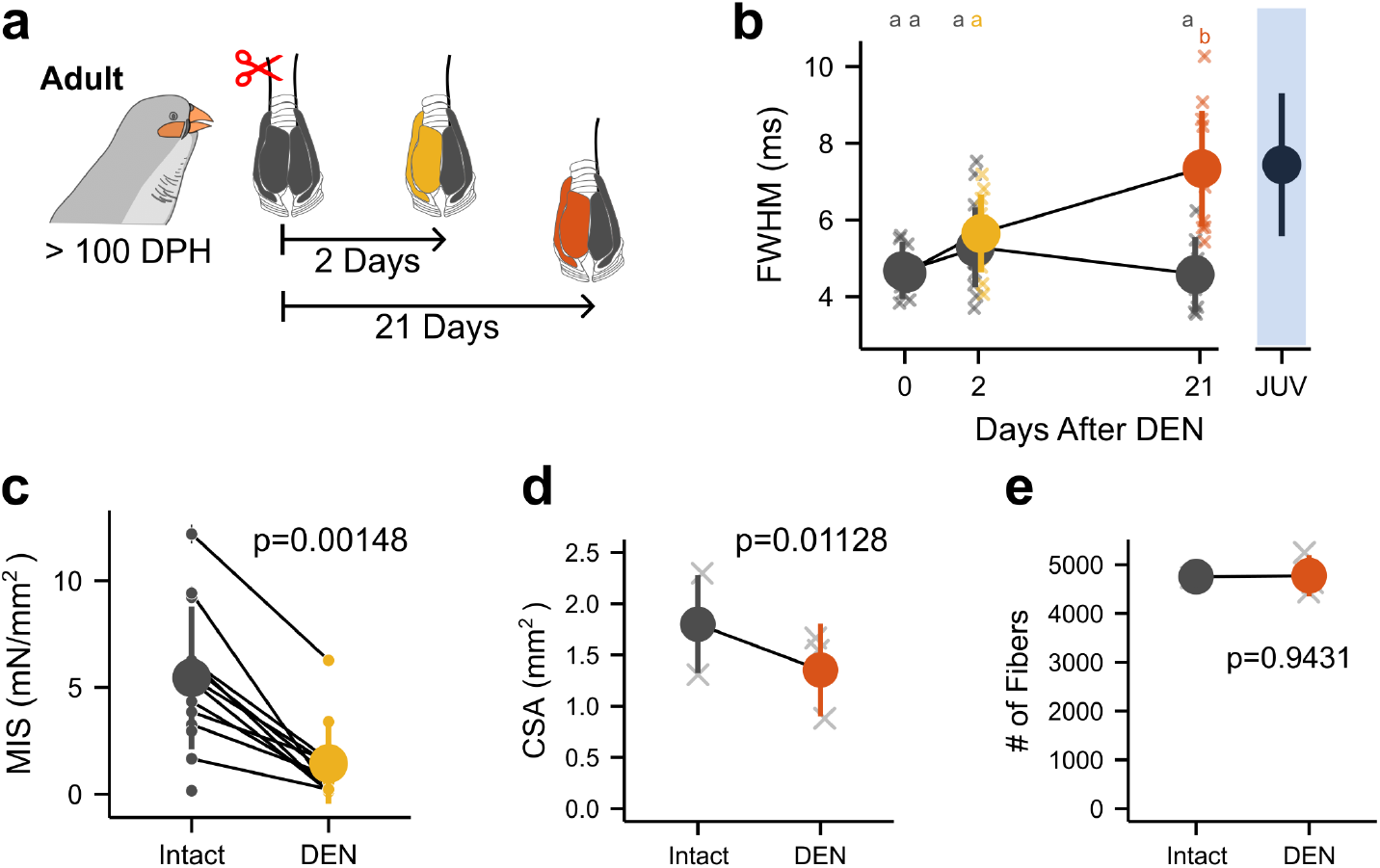
Maintaining adult peak vocal muscle performance requires daily exercise. a) Vocal muscle performance was measured 2 and 21 days after denervation (DEN) in adult males on denervated (yellow, orange) and intact control (grey) side. b) Twitch full width at half maximum (FWHM) of isometric force development increased and after 21 days reverted to juvenile speed (black). c) Maximal isometric stress (MIS) reduced fivefold two days after denervation. d) Muscle cross-sectional area (CSA) was significantly reduced 21 days after denervation, but e) muscle fibres number did not change.

## Disuse changes vocal muscle organization

Laryngeal and limb muscles contain multiple distinct fibre types with different features pertaining to force production, fatigue resistance and energy metabolisms^5^. Songbird syringeal muscles contain two fibre types^4,21^: in zebra finch males the majority (67–87%) of all muscle fibres is classified as superfast muscle (SFM) fibres^4,21^ and not immunoreactive to any available antibodies raised against heavy myosin chain (MyHC) isoforms. The remaining 13–33% of syringeal muscle fibres are smaller diameter fibres immunoreactive to an antibody recognizing mammalian fast twitch MyHCs (MY-32)^4,21^. It is currently not known if, and if so, how these two syrinx muscle fibre types respond to exercise paradigms.

To quantify if and how syringeal muscle fibre types are affected by disuse, we prevented use by unilateral denervation, stained cross-sections of the syrinx with antibodies against laminin and MY-32 (**Fig 3a**), and automatically detected muscle fibre boundaries (**Fig 3b**, see Methods) to measure their fibre CSA and MY-32 expression intensity (**Fig 3c**). On the intact side, we observed 66±3% (N=3) unstained fibres of varying size (200–1,000µm^2^) intermingled with 34%, smaller (100–400µm^2^) fibres of varying MY-32 intensity (**Fig 3a, c**), as in wild type males^4,21^. The binomial fibre area distribution combined with a unimodal MY-32 intensity distribution (**Fig 3c**) shows two fibre types, corroborating earlier results^4^. In contrast, denervation increased MY-32 intensity per fibre and resulted in a unimodal distribution of fibre area with 75% decreased CSA (**Fig 3c, d**). The total number of SFM fibres decreased (**Fig 3d**) and the SFM fraction reduced from 66±3% to 44±4%. Taken together, syringeal muscle disuse drives fibre type composition towards fibres with smaller CSA and slower contractile properties.

**Figure 3.**
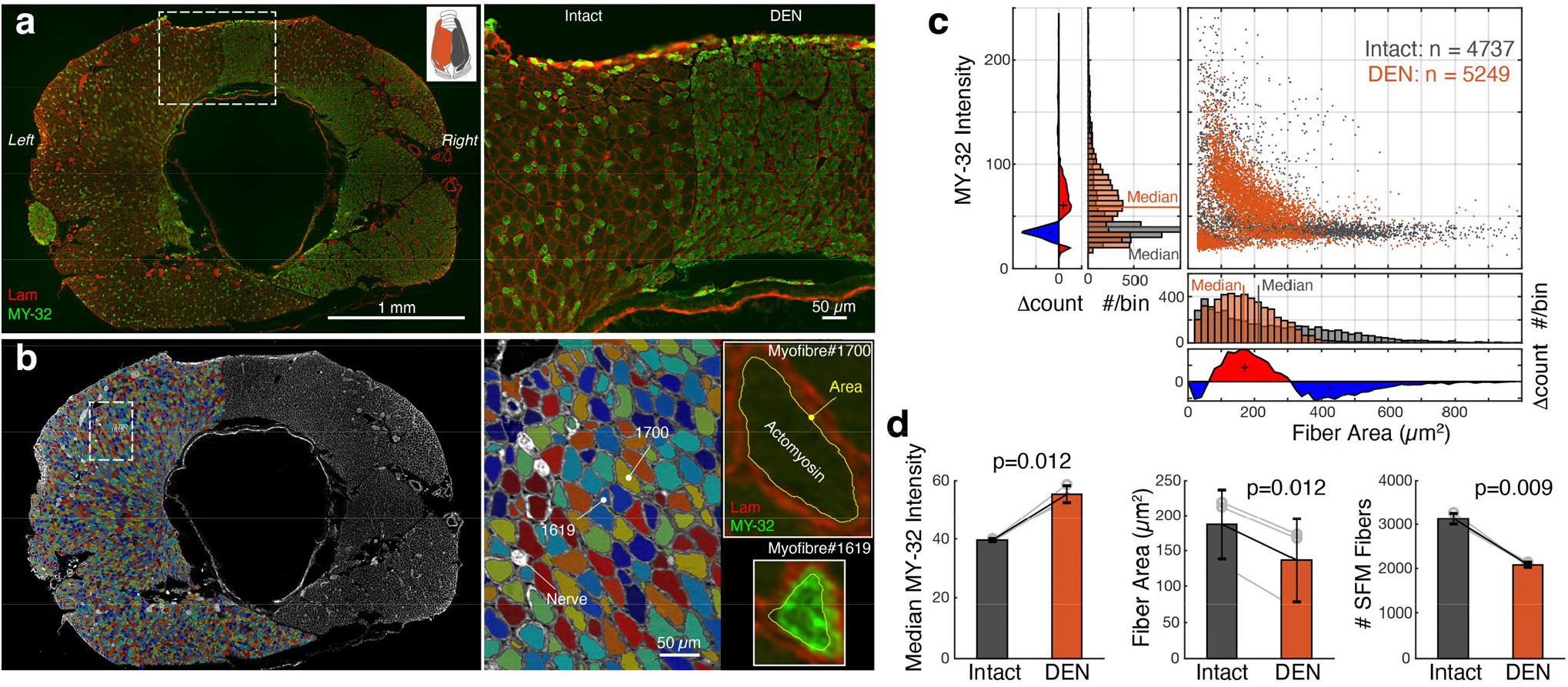
Disuse changes fibre type organization in vocal muscles. a) Cross-section (left) and detail (right) of syringeal muscle fibres of the entire adult male syrinx immunostained for laminin (Lam) and fast MyHCs (MY-32) after 21 days unilateral denervation (DEN). b) Automated myofiber detection shown in the intact hemisyrinx, faux-color-coded for fibre ID. Right, detail with (top) a typical superfast myofiber without MY-32-reactivity (myofiber #1700), and (bottom) a typical fast fibre with high MY-32 reactivity (myofiber #1619). c) After denervation (orange), fibre type composition shifts from large superfast fibres (MY-32 negative) with interspersed small fast (MY-32 positive) fibres to medium-sized fast fibres. d) Denervation increases median MY-32 reactivity, reduces median fibre area and reduces the number of SFM fibres.

## Proteomic profiling of vocal muscles

Our physiological and fibre typing data together suggest that syringeal muscle fibres respond to disuse with both quantitative and qualitative changes in protein expression^22^: reduced fibre CSA and MIS suggest reduced protein abundance, and reduced speed combined with increased reactivity to MY-32 suggest qualitative changes in expression of different MyHC isoforms. To identify and quantify disuse-induced changes in expression of proteins critical to muscle function, we performed proteomic profiling using liquid chromatography mass spectrometry first in untreated syringeal muscles (see Methods). We categorized the identified proteins according to their role in 1) force production (Sarcomere), 2) calcium handling and 3) mitochondrial function. In total we identified 450 proteins with high confidence (**Supplementary Table S2**) of which over 80% were sarcomeric, calcium handling and mitochondrial proteins (**Fig 4a, b**).

**Figure 4.**
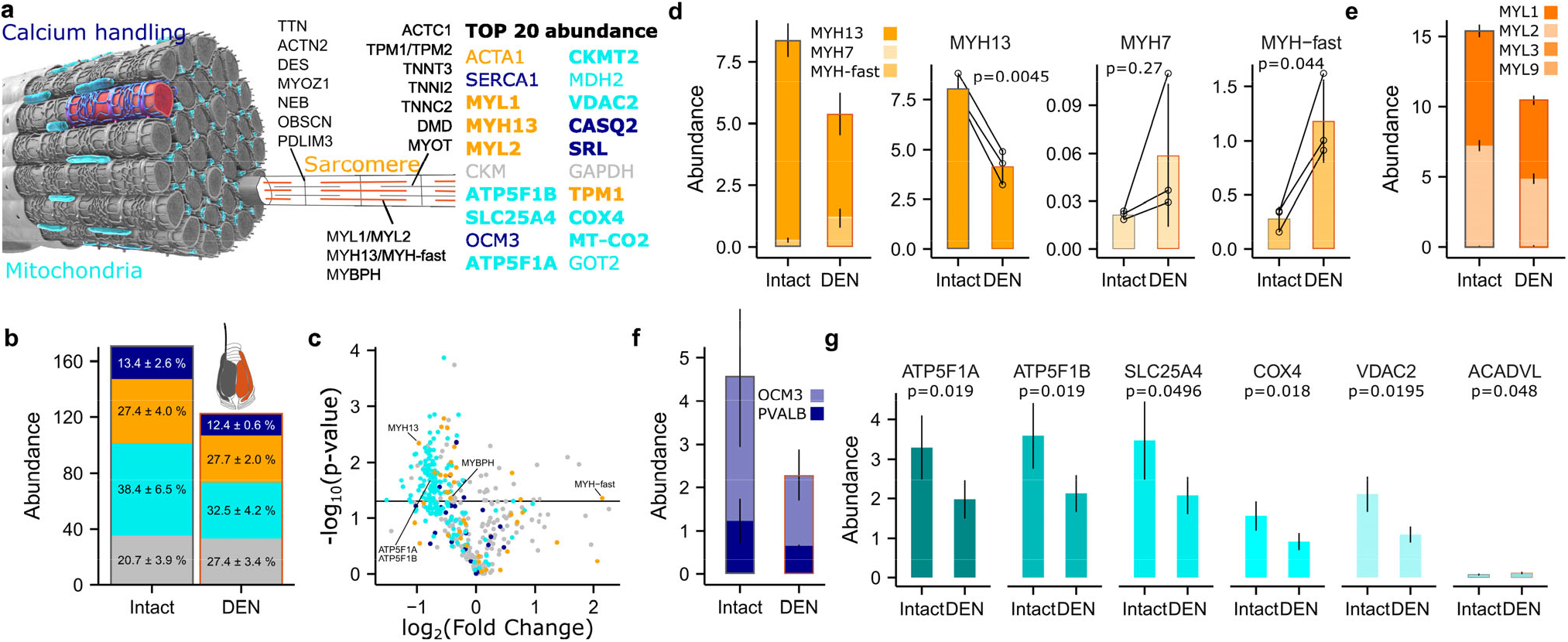
Proteomic profiling identifies sarcomeric, mitochondrial and calcium handling proteins that are downregulated by disuse. a) Schematic of superfast syringeal muscle fibre with identified proteins. The top 20 most abundant proteins are color-coded for categories sarcomere (orange), mitochondrial (cyan) and calcium handling (blue). Bold type are proteins significantly affected by denervation. b) Disuse by denervation causes downregulation of total protein abundance. c) Volcano plot of 450 identified proteins showing statistical significance over magnitude of change (FC) due to denervation. Abundance of d) total MyHC proteins, e) myosin light chains, f) parvalbumins, and g) 5 of 6 example mitochondrial proteins decrease after denervation. Relative upregulation of MYH-fast isoforms leads to a shift in MYHC composition (c).

As part of the sarcomeric proteins we detected five MyHC isoforms: MYH7 (slow), MYH13 (superfast) and 3 other MyHCs encoded by genes in the fast cluster (Uniprot accessions A0A674GPZ1, A0A674GUX9, A0A674H378). With 92±3% of the total myosin pool, MYH13 was the most abundant MyHC, conforming earlier RNA-based observations^4^, and the highest MYH13 fraction found in any muscle to date. Myosin light chains 1 and 2 (MYL1, MYL2, **Fig 4e**) and troponin subunits TNNC2, TNNI2 and TNNT3 resembled typical mammalian fast twitch striated muscle. Furthermore, calcium handling proteins involved in calcium release and reuptake (SERCA1 and RYR1) were also identical to mammalian fast twitch isoforms. Interestingly, parvalbumins - cytosolic calcium buffers - were expressed an order of magnitude higher than in mouse limb muscle^23^ **(Fig 4f)**. This provides a mechanism allowing fast muscle relaxation during bouts of muscle activity in song by pumping back temporarily sequestered cytosolic calcium back into the sarcoplasmic reticulum later between bouts of activity^3^. Mitochondrial proteins, such as proteins constituting ADP/ATP transporter (SLC25A4) and ATP-Synthase (ATP5F1A and B), were highly abundant (**Fig 4g**), making up 38±6% of the total proteome (**Fig 4b**). Taken together, the high expression of MYH13, Parvalbumins and mitochondrial proteins supports the superfast contraction speed and calcium transients as well as high energy demand of syringeal muscles.

Second, we quantified changes in protein expression after 21 days of disuse by denervation. Total protein abundance of all three functional categories reduced, except for uncategorised proteins (**Fig 4b**). Expression changed in 175 proteins, with the majority (150/175) decreasing in abundance (**Fig 4c**). Denervation affected expression of 54.4% of sarcomeric, calcium handling and mitochondrial proteins, but only 20% of uncategorised proteins (**Fig 4c**). The overall abundance of MyHCs was decreased to 64.3±1.8% of the intact side due to a decrease in MYH13 expression (96.3±1.5% to 76.5±10.0%). The expression of fast MyHC isoforms increased at the same time (MYH-fast 3.5±1.5% to 22.4±9.1%), which led to a shift in MyHC composition (**Fig 4d, e**). Because MYH13 remained highly abundant, MY-32 reactive fibres must co-express MYH13 after denervation (**Fig 3**). Expression of all calcium handling proteins, including Parvalbumins (**Fig 4f**) and all but one mitochondrial protein (ACADVL) decreased (**Fig 4c, g**). Thus, denervation reduced overall abundance of protein categories that set muscle speed and MIS, and drove composition change of MyHCs, which is consistent with our morphological and physiological data.

## Singing prevention changes muscle physiology and vocal output

Syringeal vocal muscles are active during song production, but also outside the context of singing, during calling^24^ and even rhythmically to open airways during breathing^25^. These functions may all aid to upkeep muscle performance. To test if specifically singing behaviour is required to maintain peak muscle performance in adults, we prevented adult males from singing while retaining all other functions (**Fig 5a**, See Methods). After seven days without singing, vocal muscles were significantly slower compared to control muscles of males allowed to sing freely (**Fig 5b**). Furthermore, MIS halved, but this effect was not significant (**Fig 5b**). Thus, short-term singing prevention affects speed and MIS in the same direction as short-term denervation. These data show that breathing and calling activity alone is insufficient, and adult males need regular singing exercise to maintain peak vocal muscle performance.

**Figure 5.**
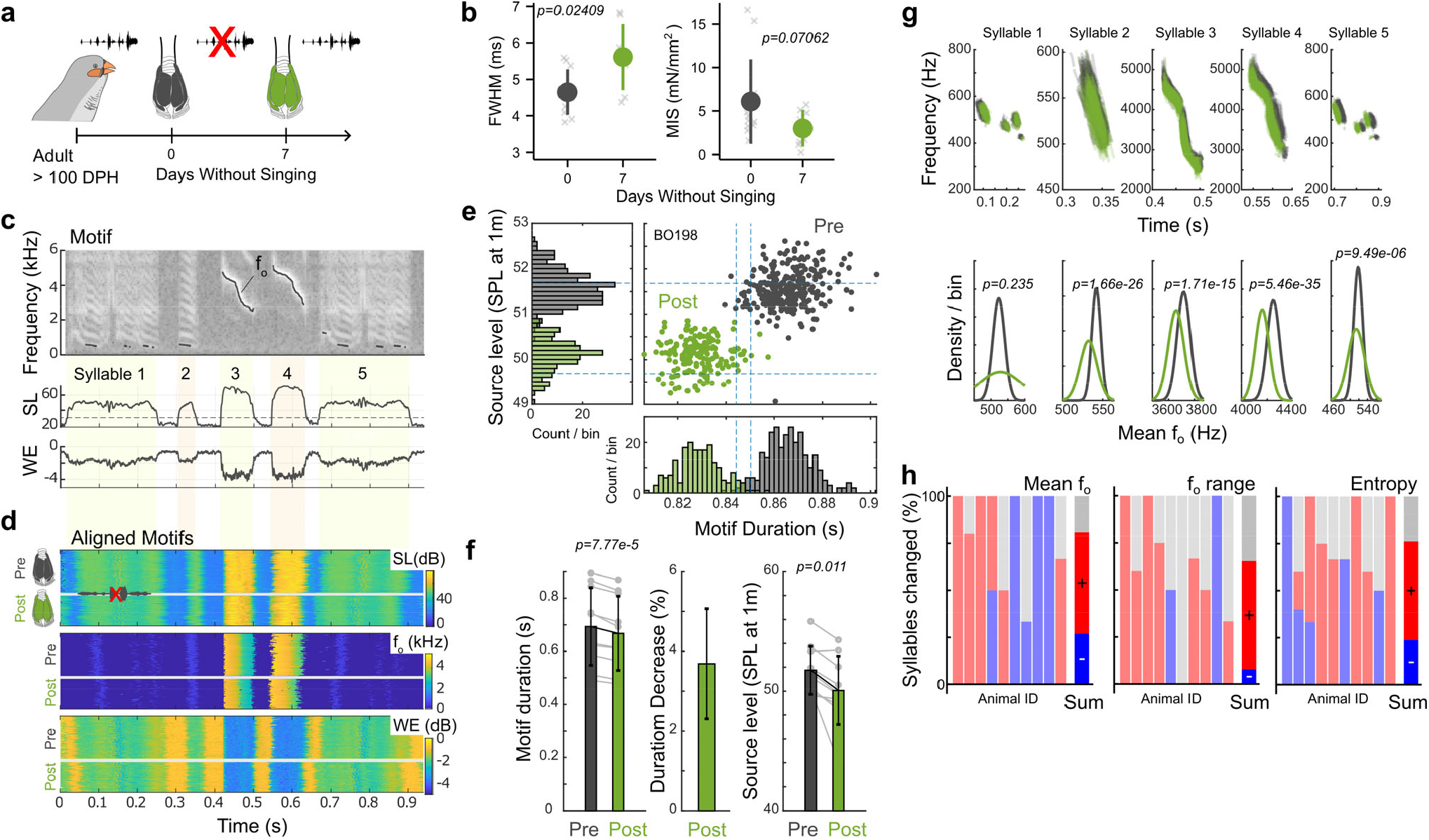
Short-term prevention of singing exercise drives changes in vocal output. a) After 7 days of singing prevention (green; See methods) b) muscle speed was significantly slower compared to singing adult males and MIS was reduced, albeit not significantly. c) Example of stereotyped song motif (individual BO198). Top to bottom: spectrogram, sound pressure, source level (SL, in dB_rms_ re. 20 µPa at 1m) and Wiener entropy (WE, in dB). Fundamental frequency (f_o_) trace is overlaid on spectrogram. d) Aligned motifs pre (n=284) and post (n=201) singing prevention of same individual color-coded for SL (top), f_o_ (middle) and WE (bottom). e) Individual BO198 and f), group data show that motif SL and duration decreased significantly due to singing prevention. g) f_o_ traces (top) and mean f_o_ distributions (bottom) per syllable of individual BO198 and h) all individuals show that singing prevention causes significant changes in f_o_, f_o_-range and WE in 81% (21/26), 65% (17/26), and 76% (32/42) of all analysed syllables of 10 animals respectively. (Red: significant increase, blue: significant decrease, grey: no statistically significant change).

To test the hypothesis that singing prevention-induced muscle changes drive acoustic changes in song, we recorded song before and after one week of targeted singing prevention, (see Methods). Vocal output is the result of interactions between neural circuitry (aka song system) and motor systems^26^. Adult zebra finches sing an individual motif consisting of several syllables. Individual syllable production and sequence is highly stereotyped (**Fig 5c, d)**. Like human speech auditory feedback corrects the motor code for deviations from the song template^9,27^. Altered acoustic feedback can drive changes in e.g. syllable pitch over days^27^, with first indications of such song system-induced corrections occurring the earliest after 6-12 hours^27^.

We analysed changes in vocal output immediately (0-2 hours) after release from singing prevention where motifs are not yet compensated by error-correction of the song production circuitry and reflect changes in the vocal periphery. Singing prevention caused several acoustic changes to song. On the motif level, duration decreased significantly from 0.69±0.15 to 0.67±0.14 s, i.e. decreasing 3.7±1.4% (N=10) (**Fig 5c - f**). Motif source level decreased from 51.7±2.0 to 50.1±2.9 dB SPL at 1m. We next extracted time-resolved f_o_ traces over individual syllables within motifs (**Fig 5g**). Although the overall shape of these f_o_ trajectories did not change, the mean f_o_ changed significantly in 81%, f_o_ range in 65% and wiener entropy in 76% of syllables from 10 animals (**Fig 5h**). Taken together, our acoustic analyses showed that already one week of targeted singing prevention causes significant shifts in vocal output due to changes in muscle performance.

There is currently insufficient insight into the mechanistic effect and function of individual syringeal muscles to link their activity to specific acoustic parameters^28^. Although vocal muscle activity can strongly correlate with an acoustic feature within a single syllable^24^ electromyographic recordings show that the strength and even polarity of correlations depends on which syllable is sung^28^. Producing specific acoustic features thus depends on the co-activation of syringeal and respiratory muscles, consistent with the notion that key acoustic parameters such as fundamental frequency are affected by several control parameters^29,30^. Our data further supports this idea, because the strength and polarity of the acoustic effect caused by reduced MIS and twitch speed of the vocal muscles are significant but also individual and syllable dependent. The reduction in motif SL caused by singing prevention is consistent with reduced subsyringeal pressure due to weaker abdominal muscles, but we did not measure physiological performance of these muscles. Motif duration is predominantly set by temperature of the motor pathway^31^ and the 3.7% duration decrease we observed here is consistent with a temperature increase of 1°C^31^. Because body temperature in birds can fluctuate up to 5°C with activity level^32^, we speculate that this temperature increase resulted from the visibly heightened state of arousal when males were allowed to sing again.

## Females prefer exercised song

Next, we tested whether the acoustic changes observed after singing prevention were perceived and meaningful to females to whom males direct song for mate attraction. Females were tested with pre or post singing songs in an established and validated operant song preference test^33^. By pecking operant keys, female zebra finches could trigger song playbacks from either pre or post singing prevention from the same male (**Fig 6a**, see Methods). Females were offered only one iteration of the recorded songs which means that to discriminate between songs, females had to detect the within motif acoustic changes, rather than variability among motifs or singing speed (parameters shown to positively affect choice^34^). Even with these cues removed, eight out of nine females showed a significant preference for a song (**Fig 6b, c**). Of the eight females, the majority (75%) preferred song from before singing prevention over song after singing prevention (**Fig 6c, d**). Females could thus perceive and distinguish the treatment-induced changes in muscle performance by listening to a single iteration of a male’s song. Importantly, they prefer songs produced by exercised vocal muscles.

**Figure 6.**
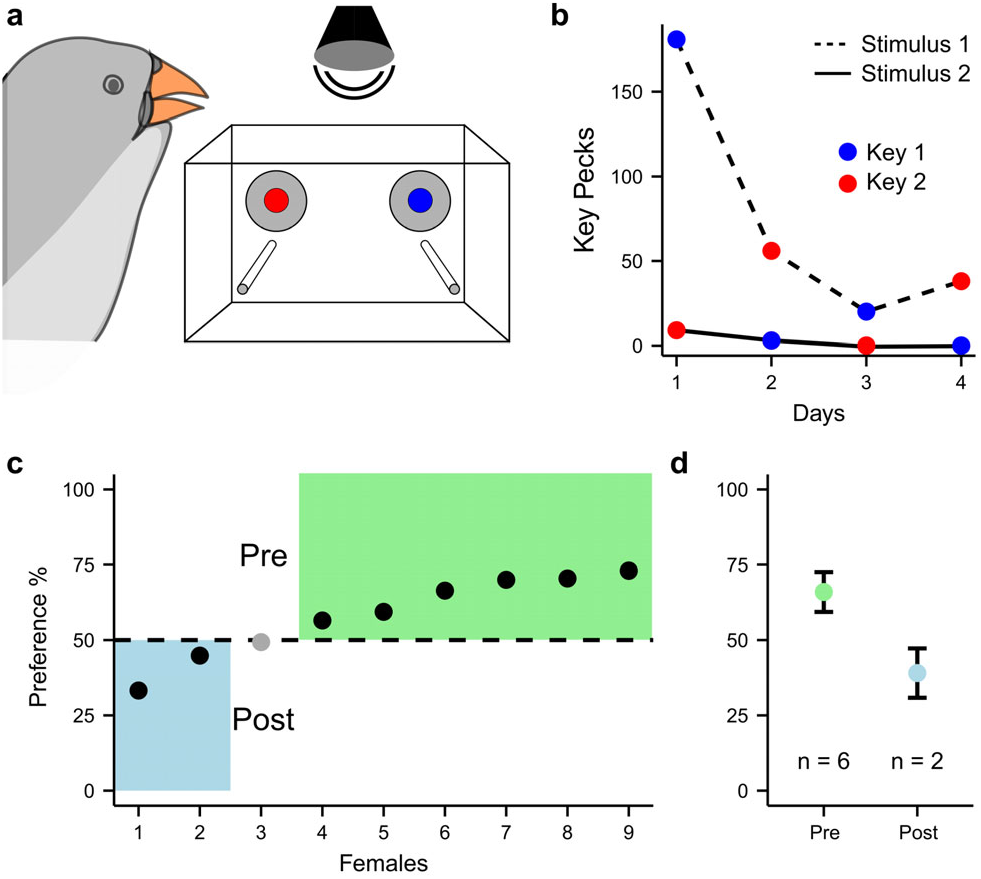
Females choose and prefer song of exercised males. a) Female choice preference test setup to elicit playback of song pre and post short-term singing prevention. b) Key peck events per day (female 6), where stimuli are side-switched daily to control for side-bias. c) Of the 9 tested females, 8 significantly preferred one stimulus above chance. Six (75 %) preferred the exercised song (pre singing prevention), d) with a mean preference of 66.0±6.62%

## Discussion

Our combined physiological, morphology, proteomics, and behavioural data show that juvenile and adult male zebra finches need to sing daily to gain and maintain peak vocal performance. Muscle exercise alters vocal output and conspecific females detect and prefer exercised songs. Conspecific receivers can thus use vocal performance as a proxy for the sender’s recent (and possibly long-term) exercise investment. The requirement of daily exercise to maintain peak performance provides a mechanistic explanation for song as an honest signal for the sender’s condition. In most vocalizing tetrapod species, muscles either precisely control the vocal organ, respiratory system and vocal tract to modulate flow-induced vibrations or directly produce acoustic signals through contractions. Because vocal and limb muscles respond to loading and unloading paradigms^2,5^, neuromuscular effects of individual exercise history on vocal output are thus likely extendable to all vocalizing vertebrates.

Our data establish that vocal exercise to optimize muscle performance is integral part of vocal skill learning in songbirds. During song learning the song system changes profoundly in size, number of neurons, connectivity and firing properties^9^. Up to the end of this period, syringeal muscles keep changing in weight, MIS and speed due to exercise^4,15,16^. Thus, both circuit remodelling^9^ and vocal muscle exercise combine to achieve the precision of final song execution and the duration of both processes contributes to and constrains the duration of extreme skill learning. What neural stimulation patterns in song specifically promote muscle hypertrophy and speed increase needs further investigation, but a possible mechanism is bursts of high frequency motor neuron firing - as observed in premotor nucleus RA^35^ - increase expression of faster MYHCs and mitochondrial content through IGF1 signalling. Interestingly, in limb muscles, first-time muscle training affects subsequent re-training duration by increasing myonuclei in existing muscle fibres^36^. These myonuclei are retained during subsequent muscle atrophy and supposedly serve as cellular muscle memory to allow faster muscle performance gains when retraining^36^. This process could explain how seasonal songbirds reach peak performance faster in their second year. Thus, early exercise can impact the speed of future skill learning with lifelong consequences.

Our data strongly suggests that song exercise is a previously unrecognized cost of adult song. Such costs are expected for sexually selected signals but direct metabolic costs seem low as oxygen consumption during singing does not elevate over silent periods^37^. Consequently, with the absence of physiological costs to produce song, costs were considered mostly developmental and song an honest indicator of past condition (the developmental stress hypothesis^38^). Our data suggest that time needed for exercise and maintaining peak performance is a previously overlooked cost to adult singing. Indeed, supplementing food to birds in the wild increases their song production^39^ suggesting a trade-off between foraging and singing. Many songbird species sing out of normally expected contexts of mate-attraction, mate-guarding and territorial maintenance, or keep singing daily - in a dawn chorus or alone - even under severely adverse conditions^10^. Song functions may thus be more diverse than previously thought^10^. The need for daily exercise to maintain peak performance may provide yet an alternative strong motivation for birds to sing daily.

Our findings suggest that the neural regulation of fibre type plasticity is equivalent between laryngeal and syringeal muscles, but different from limb muscle. In limb muscle, increasing activity increases CSA and mitochondrial function, and transforms fast into slower fibre types^40^, while unloading paradigms have the opposite effect, decreasing CSA and mitochondrial function and transforming slow into faster fibre types (**Fig 7a**). We show that syringeal muscle fibres exhibit plasticity of CSA, mitochondrial function, and fibre type by neural regulation, that are consistent with responses in laryngeal muscle (**Supplementary Table S3**). However, fibre type shifts respond oppositely to limb muscle. We propose that unloading (disuse, denervation) transforms vocal muscles from superfast to fast to slow fibre types, and vice versa in response to stimulation (song, electrical stimulation) (**Fig 7b**). Whether the opposite polarity of speed control in vocal muscles is due to neurogenic or myogenic factors remains unknown. Body size scaling considerations suggest that the fastest laryngeal fibre type expressed within a species is body size dependent^2^, and MYH13 is only present in small animals^2,41^ consistent with our data. Because normal healthy syringeal muscles seem maximally loaded^4^, we suggest loading mechanism can be best studied after experimental or natural periods of unloading, e.g., in birds singing seasonally.

**Figure 7.**
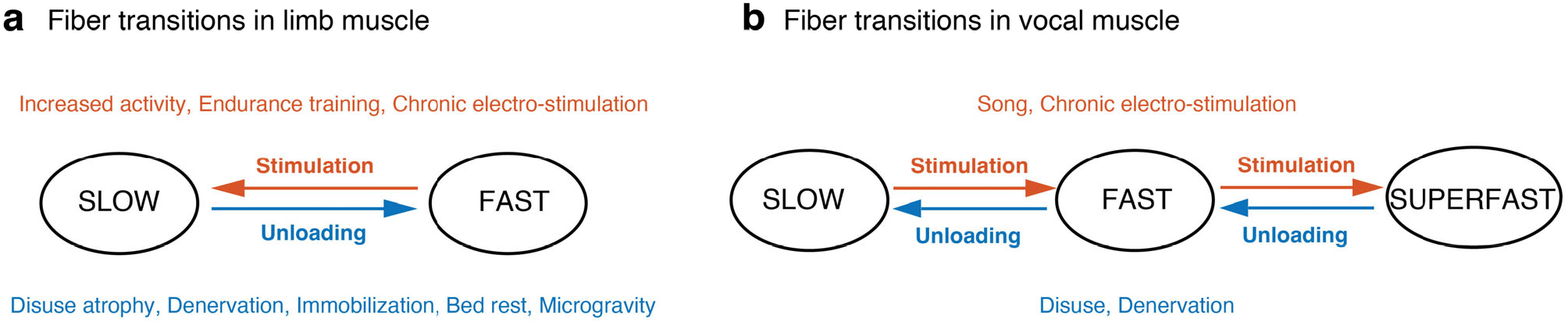
Model of neural regulation of vocal muscle fibre types. **a**, Unloading paradigms, such as disuse, denervation, bed rest etc, cause mammalian limb muscles typically to transform slow fibre types (Type 1, 2A) into fast fibre types (type 2X, 2B), while reversely training programs typically transforms fast into slower fibre types. Based on ^40^ **b**, Proposed model of fibre transitions and biomarker signatures in vocal muscle plasticity (larynx and syrinx) due to neural regulation (**Supplementary Table S3**).

Because we lack both access and experimental access to human laryngeal muscular function in vivo, there is implicit need for animal models^8^ that can be trained in paradigms similarly to human vocal training. Songbirds provide unique opportunities because 1) their song is a learned, stereotyped behaviour that is produced like human speech^9,30^, unlike rodents^42^, 2) the song motor system -from neural circuits to the vocal organ- is dedicated to song production and discrete from other vital functions and accessible to experimental manipulation^7^, which 3) enables acquisition of longitudinal datasets^19^. Songbirds furthermore 4) can be trained on complex combinatory targets for all motor systems^43^, to test coordination between motor systems which is a major pedagogical training goal of human vocal exercise^8^, 5) and provide molecular targets to model e.g. Parkinson’s disease^44^. 6). We show that syringeal muscles respond strongly to use changes in short time frames compared to larynx muscles. Therefore, we propose songbird as a powerful animal model to elucidate voice function and pathophysiology of voice disorders and to develop therapeutic interventions and pedagogical tools for voice therapy and (re)habilitation.

## Methods

### Animals

Subjects were 79 male and 13 female *z*ebra finches (*Taeniopygia guttata*). Animals at the University of Southern Denmark (SDU) were group housed in 3×3×2m aviaries on a 12:12h light:dark-cycle and provided with food, cuttlefish bone and water ad libitum. All experiments and procedures were performed in accordance with the Danish Animal Experiments Inspectorate Copenhagen, Denmark (2019-15-0201-00308).

Female preference tests were conducted at Leiden University, the Netherlands. Birds were housed in 175 × 80 × 200 cm aviaries on a 13:11h light:dark-cycle and provided with food, cuttlefish bone and water ad libitum. At least one week prior to testing animals were moved in groups of 2-4 into smaller holding cages. Experiments were approved by the committee for animal experimentation at Leiden University and the Centrale Commissie Dierproeven (CCD) of the Netherlands (1160020186606) and monitored by the Animal Welfare Body of Leiden University, in accordance with national and European legislation.

### Muscle use intervention paradigms

*Nerve cuts*. We prevented use of syringeal muscles by unilateral section of the hypoglossal nerve. Surgery was performed under a dissecting microscope. Inhalation anesthesia was induced (3%) and maintained (1%) with isoflurane (Baxter). A 10mm lateral incision was made in the skin of the neck exposing the trachea with the bilateral tracheosyringeal branches of the hypoglossal nerve (XIIts). We removed at least 5mm of the right branch by cutting the nerve with a pair of micro-scissors. The skin incision was closed with 8/0 unifilament suture (AroSurgery). Following surgery, the animals recovered in a 20×10×10 cm cage in a heated chamber (39 °C), and returned to their recording box. Nerve cuts were either performed in juvenile males before the start of song learning (30±1DPH) or in adult males (>100DPH). The juveniles were sacrificed at 65DPH and adults at 2 and 21 days post-surgery. Moreover, successful long-term denervation was supported by the disappearance of axon related proteins in the protein profiling experiments as well as immunohistochemistry for neurofilament (see below). During dissection we carefully checked for signs of reinnervation, but didn’t find evidence in any animal.

*Singing prevention*. Ten males were kept in custom-built, sound attenuated recording boxes (60×95×57cm) and vocalizations were recorded continuously45. Males were moved to recording boxes and monitored several days before entering the singing prevention paradigm to ensure they were habituated to the new environment. To prevent males from singing they were kept in the dark for 7 consecutive days, except for two to four 30-minute long feeding sessions spaced evenly throughout a day. During the feeding sessions, males were allowed to call, eat and drink ad libitum, but interrupted from singing attempts by distracting the animals. Bodyweight was monitored throughout the experiment to guarantee the animals’ wellbeing. All vocalizations were monitored continuously to ensure that birds wouldn’t start singing in the dark. On the morning of day 8 the original colony light schedule was resumed and males could sing undisturbed.

### Muscle physiology

Muscle fibre bundles were prepared and stimulated *in vitro* to record isometric force responses as previously described46. In brief, birds were sacrificed by an isoflurane overdose and the syrinx was exposed, isolated and pinned down on Sylgard-covered Petri dishes in cold, oxygenated dissection buffer47. Fibre bundles were obtained from the *m. tracheobronchialis dorsalis* (DTB). Muscle fibre bundles consisted of a subsection or the entire (denervated as juveniles and juveniles) DTB. Muscle preparations were mounted in a temperature-controlled bath perfused with oxygenated recording buffer. The rostral end of the preparation was fixed to a force transducer (Model 400A, Aurora Scientific) and the caudal end to a micromanipulator to control preparation length. After mounting, the muscle preparation was allowed to rest for 20 minutes. Muscle fibres were stimulated by a high-power follow stimulator (Model 701C; Aurora Scientific) at constant voltage using platinum electrodes. Force and stimulation signals were low-pass filtered at 10 kHz (EF120 BNC, Thor Labs) and digitized at 20 or 40 kHz (PCI-MIO-16E4, National Instruments). After optimizing the stimulation parameters at 39.0 ± 0.1 °C (as detailed in 15), we obtained seven twitch contractions and three 100 ms duration tetanic contractions at optimal length and stimulation frequency. All software to control the setup and record data was written in Matlab (MathWorks, RRID:SCR_001622). As a measure of contraction speed, we extracted the full width at half maximal (FWHM) force of single twitch stimulations, which is the time from crossing 50% force increase to decrease, and equals the previously reported t50-50 11,46. Maximal isometric stress was calculated as maximal tetanic force divide by the muscle cross-sectional area (CSA) estimated from optimal (resting) length L0, the dry weight (dry-wet conversion factor: 5) and density (1060 kg/m^3^ from ^48^) of the muscle fibres.

### Muscle morphology

Birds were sacrificed by an isoflurane overdose and immediately perfused transcardially with dissection buffer47 to remove blood cells before dissecting the syrinx on ice at SDU. Syrinxes were dried on kimwipes and frozen in an isopentane bath cooled by liquid nitrogen, and subsequently stored in cryotubes (Nunc X) at -80°C. All samples were shipped on dry-ice to Umea University for sectioning and staining.

#### Immunohistochemistry

Serial muscle cross-sections, 8-10µm thick, were cut in a cryostat (Leica CM3050S cryostat, Leica Biosystems, Nussloch Germany) at –22°C and mounted on Superfrost Plus Adhesion Microscope Slides (Menzel Gläser, Menzel GmbH & Co). We used cross-sections of the adult male zebra finch syrinx at midbody (between the most rostral part of bronchial rings B2 and 3) where all muscle fibres are present. Immunohistochemical staining was performed using well-characterized antibodies and modified standard immunohistochemical techniques. For antibody (AB) specificity and concentrations, see **Supplementary Table S4**. In brief, the sections were immersed in 5% normal non-immune goat serum (DakoX0907, Agilent Technologies Inc., CA, USA) for 15min and thereafter rinsed in 0.01M phosphate-buffered saline (PBS) for 3×5 min. The sections were then incubated with the primary Abs diluted to appropriate concentrations in PBS with bovine serum albumin in a humid environment. Incubation was carried out overnight at 4°C for mAbs M4276 and CBL212 and the next day the sections were double stained with polyclonal Ab L9393, for 1h at 37°C. After additional washes in PBS, the sections were incubated with the secondary AB (37°C for 30min) and washed in PBS 3×5 min. Bound primary ABs were visualized by indirect immunofluorescence using corresponding secondary abs, ABs prepared for multiple labeling and conjugated with fluorochrome with different emission spectra; Goat anti-Mouse IgG Alexa Fluor 488, (A-11029) and Goat anti-Rabbit IgG Alexa Fluor 568 (A-11036, Invitrogen by Thermo Fisher Scientific, Rockford, USA). The sections were thereafter washed in PBS for 3×5 min and then mounted in ProLong Gold antifade mountant or ProLong Gold antifade mountant with DAPI (4**×**,6-diamidino-2-phenylindole), for staining the nuclei (P36930, Invitrogen by Thermo Fisher Scientific, Life Technologies, Oregon, USA).

#### Morphometric analysis

All immunohistochemically stained cross-sections sections were captured using a digital camera (Leica DFC360 FX) connected to a fluorescence microscope (Leica DM6000B, Leica Microsystems CMS GmbH, Wetzlar, Germany) with a motorized table. Individual images were captured across the whole section with a 10x magnification objective to generate a high-resolution montage of the whole section. The resulting images were around 20,000 × 30,000 pixels. To separate left from right muscles, we made binary mask in Photoshop identifying left and right pixels. To significantly reduce computational time we rescaled images and mask to 10,000 pixels height.

We detected all muscle fibres per side using an automated procedure based on the laminin-stained cell borders. Analysis was implemented in Matlab (the Mathworks). We improved contrast of the laminin layer using contrast-limited adaptive histogram equalization (*adapthisteq* function) and applied the mask. We then converted the image layer first to grayscale using a global threshold 49 and second to a binary image with a 10-20% reduced shift to detect all cell borders. We removed noise areas and small negative areas (*bwareaopen* function) that were blood vessels, nerves and SR. This resulted in detection of areas between laminin containing myofibrils. To smoothen the detected myofibrillar area per muscle fibre we applied a diamond-shaped morphological structuring element to subsequently grow and shrink the areas (*imdilate* and *imerode* functions). To isolate individual muscle fibres in the binary image, we computed the distance from border to center for all areas (*bwdist* function), suppressed local mimina using the H-minima transform50 (*imhmin* function) and computed the watershed segmentation (*watershed* function). The segmentation accurately detected the myofibrillar area of the muscles, but thus left out a large area in between fibres containing other cellular components. To estimate the total muscle CSA per hemisyrinx we therefore dilated all fibres with a diamond-shaped morphological structuring element (radius 10) to fill up the space in between fibres.

For each resulting fibre area we computed multiple morphological features (area, perimeter, eccentricity, solidity, major and minor ellipse length and ratio, circularity) and used feature constraints to remove false detections. We used the detected areas to measure mean MY-32 expression intensity per fibre on the MY-32 image layer. To quantify the fibre area and MY-32 expression distribution we calculated histograms with fixed bin widths. To quantify changes in these distributions due to experimental interventions, we calculated probability density function estimates to correct for differences in the total number of fibres in each dataset (*histcounts* function, ‘*pdf’* normalization). We considered fast fibres to have a mean MY-32 expression intensity >50, and superfast muscle (SFM) fibres a mean MY-32 expression intensity ≤50. This boundary clearly separated the unstained from stained fibres in the untreated condition. The obtained fractions of 66% SFM fibres correspond well with the 67% extracted from Mead et al.4 who used a Kolmogorov-Smirnov optimization to separate two populations of fibres in a single syringeal muscle (DTB), and are lower than the 87% reported in Christensen et al.21 who manually counted fibres of the entire syrinx.

### Proteomics

To determine the relative abundance of muscle proteins in the DTB, 0.6-1.2 mg samples were shipped on dry ice to the University of Vermont and analysed by label-free proteomic analyses as previously described51. Briefly, the muscle samples were solubilized in Ragigest SF Surfactant (Water), reduced, alkylated, and digested with trypsin (Promega). The resultant peptides were separated by ultra-high pressure liquid chromatography and directly infused into a Q Exactive Hybrid Quadrupole-Orbitrap Mass Spectrometer (Thermo Fisher Scientific). Data were collected in data dependent mode and recorded in .raw files. Peptides were identified and liquid chromatography (LC) peak areas were determined using Proteome Discoverer 2.2 to search against the zebra finch (*Taeniopygia guttata*) database downloaded from UniProt (11/22/2022). The searches accounted for the presence the following posttranslational modifications: oxidation (M, P), dioxidation (M), phosphorylation (S, T, Y), carbamidomethyl (C), and acetylation (N-terminus of protein). The LC peak areas were exported into Excel and the abundance of the top 3 ionizing peptides from each protein isoform of interest were used to quantify protein abundances. Results were manually curated to remove redundant peaks and gene names were curated to follow the HGNC nomenclature52. After removing redundant entries, LC peaks were normalized to the total sum of peaks and Histone H4 (Uniprot accession: B5FXC8) (Supplemental table S2). Expression of all MyHC genes belonging to the fast/developmental cluster on chromosome 18 (NC_044230.2 5,745,795-6,139,115), except MYH13 were quantified together using LC peaks from shared peptides and the protein is called MYH-fast in the entire MS. This strategy was chosen as not all gene models were identifiable by unique peptides. Proteins were grouped into 4 groups: sarcomeric, Ca-handling, mitochondria, other based on their subcellular localization and function. Information on subcellular localization and or function of proteins was extracted from Uniprot (http://www.uniprot.org/), gene cards (https://www.genecards.org/) and literature searches.

### Song recording and analysis

In each recording box, sound was monitored continuously by Sound Analysis Pro45 and recorded by an omnidirectional microphone (Behringer ECM8000) mounted 12 cm above the cage, digitized at 16-bit and 44.1kHz (Roland octa capture, amplification 40 dB). Recording chain sensitivity was calibrated with a 1 Pa, 1 kHz tone (sound calibrator model 42AB, G.R.A.S., Denmark).

Per animal (N=10 males) we defined a sound segment containing the motif and used cross-correlation to detect and isolate motif segments in all pre and post song files. We recorded 976±455 (median: 881, range: 554-2087) and 535±288 (median: 581, range: 82–934) motifs pre and post singing prevention respectively. We bandpass filtered the sound between 200-12,000 Hz using a 2nd order Butterworth filter with zero-phase shift implementation (*filtfilt* function). We divided the signal into 4 ms duration bins with 0.5 ms steps and calculated the following acoustic features per bin: aperiodicity, power, source level, and f_o_ using the Yin algorithm53. We aligned the motifs to the highest correlation of the source level (*finddelay* and *circshift* function) for the pre and post singing prevention separately. To ensure we compared acoustic feature trajectories of motifs with the same syntax, we focused on the most common syntax and omitted all other motifs.

We used a fixed SL threshold per individual to segment sound and silence within the motif, and removed segments of sound and silence below 30 ms and 40 ms duration, respectively. The remaining binary signal represented the syllables and was used to extract syllable on- and offset timing, syllable and gap duration. Motif duration was the summed duration of all detected syllables and gaps.

Next, we calculated the mean, minimal, maximal and range of f_o_ per syllable, after removing spurious f_o_ detections within syllables by removing jumps between adjacent bins over 100 Hz and fo trajectories below a duration of 5 ms. We omitted syllables where pitch detection was not robust. In total, we analysed 485±261 (median: 393, range: 233-1093) and 296±157 (median: 249, range: 81–532) motif iterations pre and post singing prevention.

### Female choice experiment

(for more details also refer to Supplementary Methods) For the song preference test, we used a previously validated operant paradigm33. Nine female zebra finches were moved into one of nine identical experimental setups consisting of a wire-mesh cage (70 × 30 × 45 cm) with a solid back panel with operant keys, each placed in separate sound-attenuated chamber. From the first (left) and fifth (right) of five equally spaced perches, a bird could peck a 5 cm diameter white piezoelectric circular plate (response key) with a small embedded red light-emitting diode LED (5 mm diameter) at the top. When pecked a custom-built minicomputer (sound chip Oki MSM6388, Tokyo, Japan) and laptop (Sony Vaio E series, Sony, Minato) placed outside the experimental chamber registered the activation time of the response key and triggered an acoustic playback of an assigned stimulus via a loudspeaker (Vifa 10BGS119/8, Viborg) suspended from the ceiling at 1 m above the cage. Stimuli were played back at 70 dB re 20 µPa at the central perch (Voltcraft sl451 sound level meter, fast response setting, A-weighting). During all training and testing, experimenters were blinded with respect to whether a stimulus was recorded pre- or post-treatment.

*Motif selection and stimulus construction:* Because both total number of syllables and source level are known to influence preference54,55 we ensured that stimuli pre and post were equally long and loud. From all pre and post recorded songs, we first identified motifs that were within 4 ms duration and 1-2 dB SL of the mean motif duration and mean SL pre and post singing prevention (**Fig 5c**). In one individual the pre and post SL differed so much we used a 30ms and 6 dB range to find overlap. From the overlapping motifs, we randomly picked one pre and one post motif per male and used this to construct a natural song bout for each male. We exchanged the motifs in a bout sung by that male before singing prevention to retain a natural bout organization including introductory notes.

*Preference analysis*. The pecking events and cumulative learning curves of each female were checked daily and preference testing began the day after the initial pecking had changed from incidental pecking to an exponential increase of frequent operant responses on both sides34. Each test lasted 4 days and each night we switched the assignment of the stimulus songs between left and right keys to control for side preferences. We calculated female preference as the sum of keypecks for one stimulus over 4 days divided by the sum of total keypecks. Individual female preferences were analysed with G tests against 50% chance level using Williams’ correction.

### Statistics

No formal methods to predetermine sample size were used; sample sizes are similar to those used in the field. Randomization and blinding was performed when analysis wasn’t automated. Details are described in each section of the methods. Statistical results and setting for all tests applied are presented in Supplementary Table S1. Data are presented as mean values ± 1 S.D throughout the manuscript.

## Data availability

Data for figures is available upon request from authors.

## Code availability

Code is available on request from authors.

## Acknowledgements

Peter Snelderwaard, Maria Anthonsen, Bianca Jorgensen and Anna-Karin Olofsson for technical support. Sonja Jacobsen, Emilie Radoor and Emilie Jensen for animal care.

## Author Contributions

IA and CPHE designed experiments. IA carried out the muscle physiology experiments and analyses. PS carried out the fibre typing experiments, CPHE carried out the image analyses. NW, MP conducted the proteomic data and MP and IA analysed the proteomics dataset. IA carried out the acoustic recordings and CPHE analysed the data. IA, KR and CPHE conducted the female choice experiments and IA carried out the analysis. IA and CPHE wrote the first draft of the manuscript, all authors contributed to the final draft.

## Additional information

Supplementary Information is available for this paper.

