## Supplemental information for "Peak performance singing requires daily vocal exercise in songbirds"

### Supplementary Information

#### Table S1. Summary of statistics

See enclosed Excel table.

#### Table S2. Proteomics: LC peaks of intact and denervated DTB muscle

See enclosed Excel table.

**Table S3. Fiber plasticity in laryngeal and syringeal vocal muscle features due to neural modulation.** Changes in MyHC composition in loading paradigms gives a mixed image: Chronic electrostimulations shows no detectable shift or a slight increase in slower MyHCs, while cross-innervation with a faster nerve leads to an increase of faster MyHCs, reinnervation leads to slower MyHC composition. In contrary to denervation and unloading studies in general, where neural drive is inevitably decreased, information about the introduced change in neural drive is lacking in loading studies.

| **Unloading** | **Larynx** | **Evidence** | **Syrinx** | **Evidence** |
|---|---|---|---|---|
| CSA | ↓ | References 1-8 | ↓ | This study |
| MyHC isoform/speed | Fast isoform ↓ Slow isoform | References 1,2,4,9,10 but see 11 | Fast ↓ Slow | This study |
| Oxidative potential Mitochondria | ↓ | Reference 3 | ↓ | This study |
| **Loading** | **Larynx** | | **Syrinx** | |
| CSA | ↑ | References 12,13 but see 14 | (↑) | Increase after onset of song learning; Reference 15 |
| MyHC isoform/speed | ↑ | Reference 16 but see 12,14 | (↑) | Increased MYH13 expression after onset of song learning; References 17,18 |
| Oxidative potential Mitochondria | ↑ | Reference 14 | (↑) | Unusually high mitochondrial VPE and cristae density in freely singing animals; References 17,19 |

**Table S4. Primary antibodies used for immunohistochemistry**

| Antibody | Target | Gene (human) | Dilution | Clone | Source |
|---|---|---|---|---|---|
| **M4276** | Fast MyHC | *MYH1, MYH2* | 1:500 | MY-32 | www.sigmaaldrich.com/DK/en/product/sigma/m4276 |
| **L9393** | Laminin | *LAMA1* | 1:500 | Polyclonal | www.sigmaaldrich.com/DK/en/product/sigma/l9393 |
| **CBL212** | Neurofilament | *NEFH* | 1:500 | RT97 | www.sigmaaldrich.com/DK/en/product/mm/cbl212 |

*Official gene nomenclature according to HGNC. (https://www.genenames.org/)

### Supplemental Methods

#### Female preference testing

Birds in Leiden, the Netherlands, were housed and tested in climate regulated rooms (19–22 °C and 40–60% humidity) of the bird facility. Lights were on from 0700 to 2030 h (starting and ending with a 15 min twilight phase). Seed mix was (Deli Nature 56-Foreign finches super, Schoten, Belgium) enriched with minerals and vitamins (GistoCal, Raalte, the Netherlands) and supplemented once per week with egg food. The majority of females had breeding experience and all had been housed with males at least in hearing distance.

For the song preference test, wire mesh cages were each placed in separate sound-attenuated chambers (height: 250 cm, width × length irregular quadrilateral minimally 106 × 158 × 304 × 387 cm). Tests started by moving birds individually from their group cage into the operant chamber.

For training, females were first left to explore the cage with the operant set up on active (red LEDs and playback reward switched on during lights on) as some females quickly discover the link between key pecking and song reward by autoshaping. Females that did not start key pecking spontaneously within the first day were given training sessions twice daily for 20 min until operant responses were logged. During shaping, the experimenters (K.R., I.A., C.P.H.E., outside the test chamber behind a one-way mirror) initially drew females' attention to the keys by flashing the LED lights before playing the song reward and then gradually rewarding all behavior leading to closer approach and exploration of the keys, taking care to reinforce the keys on both sides (details on training, see [20]). During initial training (involving 10 females and 10 stimulus sets), songs were randomly chosen from the pool of pre and post singing prevention to see whether females were motivated to peck for either stimulus category. During this first testing round 7/10 females learned to peck the keys and were highly motivated to hear songs (>30 key pecks/day) and as previously reported for this species females preferred longer songs[20,21]. For the second, actual preference test all successfully trained females plus 3 additional females were tested with the actual test songs (See stimulus preparation main text).
